# Systematic phenomics analysis of ASD-associated genes reveals shared functions and parallel networks underlying reversible impairments in habituation learning

**DOI:** 10.1101/687194

**Authors:** Troy A. McDiarmid, Manuel Belmadani, Joseph Liang, Fabian Meili, Eleanor A. Mathews, Gregory P. Mullen, James B. Rand, Kota Mizumoto, Kurt Haas, Paul Pavlidis, Catharine H. Rankin

**Affiliations:** Djavad Mowafaghian Centre for Brain Health, University of British Columbia, 2211 Wesbrook Mall, Vancouver, British Columbia V6T 2B5, Canada; Department of Psychiatry, University of British Columbia, 2255 Wesbrook Mall, Vancouver, British Columbia V6T 2A1, Canada.; Michael Smith Laboratories, University of British Columbia, 2185 East Mall, Vancouver, British Columbia V6T 1Z4, Canada; Genetic Models of Disease Research Program, Oklahoma Medical Research Foundation, Oklahoma City, Oklahoma 73104; Oklahoma Center for Neuroscience, University of Oklahoma Health Sciences Center, Oklahoma City, Oklahoma 73104; Department of Zoology, University of British Columbia, 2350 Health Sciences Mall, Vancouver, British Columbia V6T 1Z4, Canada; Department of Psychology, University of British Columbia, 2136 West Mall, Vancouver, British Columbia V6T 1Z4, Canada

**Keywords:** Autism Spectrum Disorder, Neurodevelopmental disorders, *Caenorhabditis elegans*, Phenomics, Behavior, Habituation learning, Genetic networks, CRISPR-Cas9, Neuroligin, Variants of uncertain significance

## Abstract

A major challenge facing the genetics of Autism Spectrum Disorders (ASD) is the large and growing number of candidate risk genes and gene variants of unknown functional significance. Here, we used Caenorhabditis elegans to systematically functionally characterize ASD-associated genes in vivo. Using our custom machine vision system we quantified 26 phenotypes spanning morphology, locomotion, tactile sensitivity, and habituation learning in 87 strains each carrying a mutation in an ortholog of an ASD-associated gene. We identified hundreds of novel genotype-phenotype relationships ranging from severe developmental delays and uncoordinated movement to subtle deficits in sensory and learning behaviors. We clustered genes by similarity in phenomic profiles and used epistasis analysis to discover parallel networks centered on CHD8•chd-7 and NLGN3•nlg-1 that underlie mechanosensory hyper-responsivity and impaired habituation learning. We then leveraged our data for in vivo functional assays to gauge missense variant effect. Expression of wild-type NLG-1 in nlg-1 mutant C. elegans rescued their sensory and learning impairments. Testing the rescuing ability of all conserved ASD-associated neuroligin variants revealed varied partial loss-of-function despite proper subcellular localization. Finally, we used CRISPR-Cas9 auxin inducible degradation to determine that phenotypic abnormalities caused by developmental loss of NLG-1 can be reversed by adult expression. This work charts the phenotypic landscape of ASD-associated genes, offers novel in vivo variant functional assays, and potential therapeutic targets for ASD.

## INTRODUCTION

Autism Spectrum Disorders (ASD) encompass a clinically and genetically heterogeneous group of neurodevelopmental disorders characterized by diverse deficits in social communication and interaction, restrictive repetitive behaviors, and profound sensory processing abnormalities.^1–5^ The fifth edition of the Diagnostic and Statistical Manual of Mental disorders combines autistic disorder, Asperger disorder, childhood disintegrative disorder and pervasive developmental disorder not otherwise specified into the single grouping of Autism Spectrum Disorder.^1^ Despite extensive study, there is currently no unanimously agreed upon structural or functional neuropathology common to all individuals with ASD and there is little understanding of the biological mechanisms that cause ASD.^3^ The most promising avenue for research into ASD has stemmed from the observation that they have a strong genetic component, with monozygotic concordance estimates of ∼70-90% and several distinct highly penetrant genetic syndromes.^3,6–14^

Rapid advances in copy number variation association, whole-exome, and more recently, whole-genome sequencing technology and the establishment of large sequencing consortia have dramatically increased the pace of gene discovery in ASD.^15–28^ There are now >100 diverse genes with established ties to ASD, many of which are already being used in diagnosis. Importantly, each of the genes individually account for <1% of cases.^3,4,25,29–31^ Some of these genes have fallen into an encouragingly small set of broadly defined biological processes such as regulation of gene expression (e.g. chromatin modification) and synaptic neuronal communication, and have begun to offer valuable insights into the biological mechanisms underlying this heterogeneous group of disorders.^3,15,22,24,25^ However, thousands of additional mutations in these and many other genes have been identified in individuals with ASD and their roles as causative agents, or their pathogenicity, remains ambiguous. Indeed, our ability to sequence genomes has vastly surpassed our ability to interpret the genetic variation we discover. Thus, there are two major challenges facing the genetics of ASD: 1) the large, and rapidly growing number of candidate risk genes with poorly characterized biological functions and; 2) the inability to predict the functional consequences of the large number of rare missense variants. Difficulties in rare missense variant interpretation stem in part from constraints on computational variant effect prediction and a paucity of *in vivo* experimental variant functional assays.^32–35^ This lag between gene discovery and functional characterization is even more pronounced when assessing the role of putative ASD risk genes and variants in complex sensory and learning behaviors. Beyond limited gene level functional information there is even less known about how ASD-associated genes functionally interact in networks. As such, there is a great need to rapidly determine the functions of ASD-associated genes, the functional consequences of variants of uncertain significance, and delineate complex functional genetic networks among ASD-associated genes *in vivo*.

The genetic model organism *Caenorhabditis elegans* is a powerful system for the functional analysis of disease-associated genetic variation, particularly for high-throughput *in vivo* characterization of risk genes identified through genomics.^36^ *C. elegans*’ fully sequenced and thoroughly annotated genome as well as its complete connectome have fueled numerous human disease discoveries, including the role of the insulin signaling pathway in normal and pathological aging and the identification of presenilins as part of the gamma secretase complex.^36–43^ There are clear *C. elegans* orthologs for >50% of human genes, and human genes have repeatedly been shown to be so structurally and functionally conserved that they can directly replace their *C. elegans* ortholog.^36,44–49^ There are well-annotated libraries of *C. elegans* strains with deletion alleles available for >16,000 of the ∼20,000 protein coding genes.^50–52^ *C. elegans* small size and rapid hermaphroditic mode of reproduction (3 days from egg to egg-laying adult) allows for the routine cultivation of large numbers of isogenic animals. Further, CRISPR-Cas9 genome engineering has proven to be reliable and efficient in *C. elegans*, and unlike most organisms analyzed to date, multiple rigorous whole-genome sequencing studies have revealed no evidence of significant off-target effects due to CRISPR-Cas9 genome editing in this organism.^44,53–59^ Finally, we developed the Multi-Worm Tracker (MWT), a machine vision system that allows for comprehensive phenotypic analysis of large populations of freely behaving animals while they perform complex sensory and learning behaviors.^60,61^ Multiplexing by running several trackers in parallel allows for analysis of multiple measures of morphology, locomotion, mechanosensory sensitivity, as well as several forms of learning in hundreds of animals simultaneously.^62–64^

Here, we developed a scalable phenomic characterization pipeline designed to discover the functions of ASD-associated genes by systematically inactivating each gene in a model organism and observing the phenotypic consequences using machine vision (**Figure 1A**). We present data summarizing scores on 26 quantitative phenotypes spanning morphology, baseline locomotion, tactile sensitivity, and habituation learning in 87 strains of *C. elegans* each carrying a mutation in an ortholog of an ASD-associated gene, revealing hundreds of shared and unique genotype-phenotype relationships. We clustered strains based on phenotypic similarity to discover novel functional interactions among ASD-associated genes, which we validate with epistasis analysis. Further, we use the novel phenotypes for *in vivo* functional assays to assess missense variant effect, and to determine whether phenotypic alterations are reversible using targeted protein degradation methods based on degrons.

**Figure 1.**
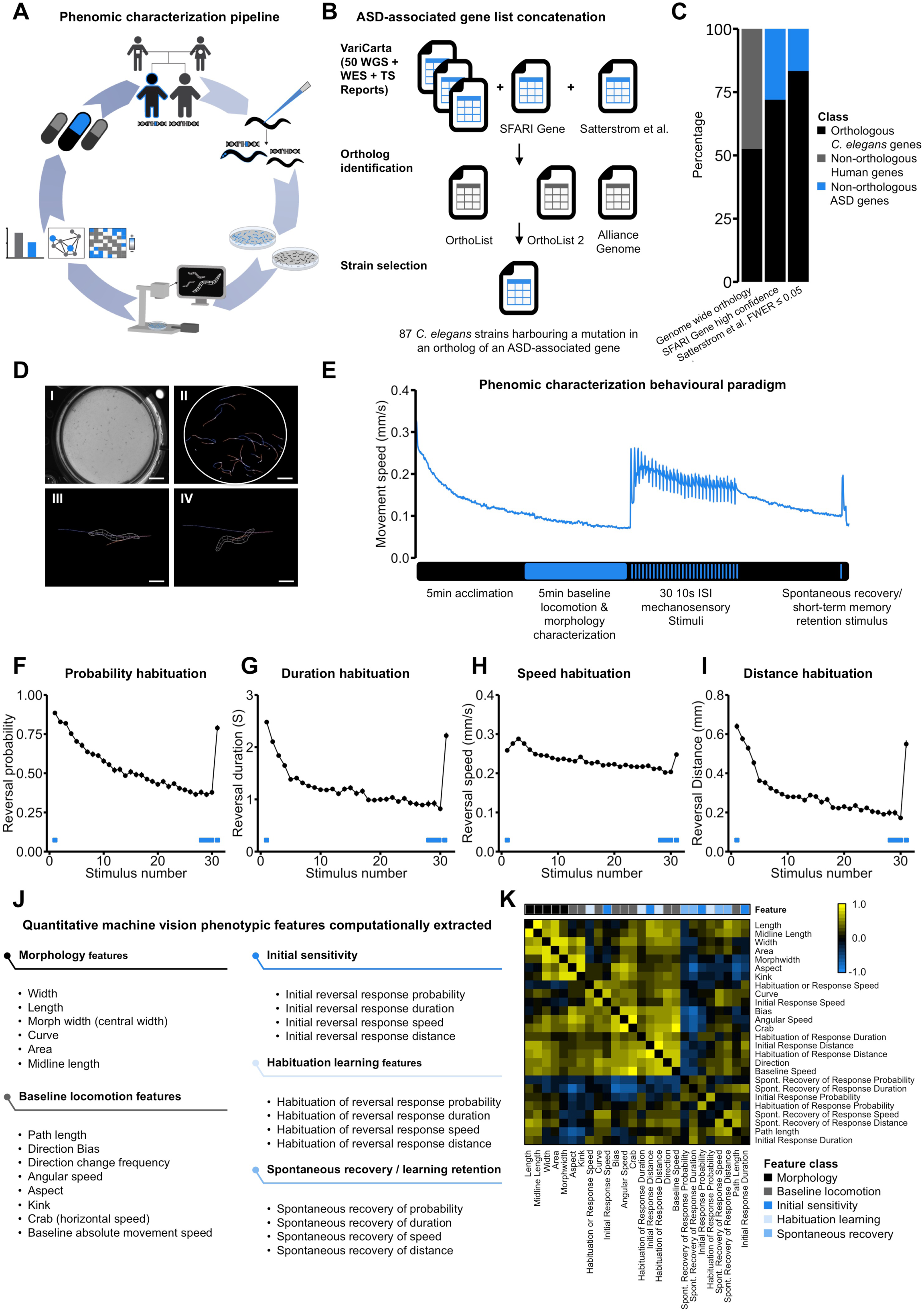
A scalable ASD-associated gene ortholog identification and phenomic characterization pipeline. **A)** Schematic illustration of the pipeline: Animals carrying mutations in orthologs of ASD-associated genes are systematically genetically engineered using CRISPR-Cas9 or ordered from the Caenorhabditis Genetics Centre (CGC), large synchronous isogenic populations of wild-type and mutant animals are grown and their morphological, baseline locomotion, initial sensitivity, habituation learning, and memory retention phenotypes are characterized using The Multi Worm Tracker. Novel genotype-to-phenotype relationships derived from the machine vision data are then used to cluster strains based on phenotypic similarity, establish variant functional assays, and test reversibility of phenotypic alterations. **B)** Schematic of databases and tools used during putative risk gene list concatenation and ortholog identification. Whole-Genome Sequencing (WGS), Whole-Exome Sequencing (WES), Targeted Sequencing (TS). **C)** Orthology between human genes and *C. elegans* genes. A greater proportion of high-confidence ASD-associated genes have *C. elegans* orthologs (blue, 72% 18/25 SFARI Gene, 83% 20/24 Satterstrom et al., 2018) compared to all human genes (gray, 53% 10,678/23,010). **D)** The Multi-Worm Tracker delivers stimuli and performs image acquisition, object selection, and parameter selection in real time while choreography software extracts detailed phenotypic information offline (panels) I) petri plate of *C. elegans* II) A petri plate of *C. elegans* selected for analysis by the Multi-Worm Tracker III) A Multi-worm tracker digital representation showing the degree of phenotypic detail. An example behavior scored by the Multi-Worm Tracker: the *C. elegans* response to a mechanosensory tap to the side of the Petri plate is brief backwards locomotion (from III to IV). Scale bars are 1cm, 1cm, 0.25mm, 0.25mm from I-IV. **E)** Phenomic characterization behavioral paradigm plotted alongside a single phenotype, absolute movement speed. Following a 5min acclimation phase a further 5min period is recorded from which multiple measures of morphology and baseline locomotion are extracted. Beginning at 10min 30 mechanosensory stimuli are delivered at a 10s ISI to which the animals initially reverse and then habituate, allowing for assessment of multiple measures of initial sensitivity and learning. The Habituation phase is followed by a 5min recovery period before administering a 31st stimulus to assess spontaneous recovery from habituation/short-term memory retention. **F-I)** Multiple measures of habituation learning of the same reversal responses exhibit different extents and rates of learned response decrement. Each point represents a reversal response. Data are shown as mean±s.e.m using the number of plates as n. **J)** List of quantitative machine vision phenotypic features computationally extracted and their predefined subclasses. **K)** Hierarchically clustered correlation matrix illustrating varied moderate correlations between features. Pearson’s r is shown.

## RESULTS

### ASD-associated genes are highly conserved to *C. elegans*

We set out to systematically characterize the phenomic profiles of *C. elegans* strains carrying inactivating mutations in orthologs of the ASD-associated genes with the highest number of rare likely gene-disrupting and missense variants reported in the literature. We identified *C. elegans* orthologs of the top 50 ASD-associated genes by variant count according to our ongoing meta-analysis of rare variants, Varicarta (https://varicarta.msl.ubc.ca/)^65^. We also identified a large number of orthologs from SFARI gene (https://gene.sfari.org/) “Syndromic” and “High confidence” categories.^66^ Of note, 72% (18/25) of genes listed in the High confidence category of SFARI gene and 83% (20/24) of genes most confidently associated with ASD by Satterstrom et al. (2018) have a clear *C. elegans* ortholog according to OrthoList2, a compendium of human-*C. elegans* orthology prediction algorithm results.^46^ This is substantially higher than the 53% estimated genome wide orthology between humans and *C. elegans* (10,678/23,010)^46^ suggesting an exceptionally high conservation of biological processes core to the pathology of ASD (**Figure 1B,C**).

Rapid advances in gene discovery and orthology prediction have altered the gene lists used during the course of this project, a challenge facing similar recent systematic investigations of ASD-associated gene function.^67,68^ Despite these shifts, our list still covers 82% (14/17) of the most strongly associated genes from Satterstrom et al. and 87% (13/15) of the SFARI Gene high confidence category listed genes (**Table S1**) with a viable ortholog deletion or severe missense allele available. This led to a mix of currently defined high- and mid-confidence ASD-associated genes, giving us a unique opportunity to study putative ASD-associated genes of completely unknown function alongside well established genes with known roles in neurodevelopment and sensory processing (See **Table S1** for a complete listing of characterized strains, orthology relationships, and ASD-association confidence).

We then used our machine vision MWT (**Figure 1D**) to systematically characterize the 87 *C. elegans* strains with a mutation in an ortholog of an ASD-associated gene (strains include 66 with mutations in different genes and 21 strains with secondary alleles to a subset of these genes, see **Table S1** and **Methods**). As it was unknown how perturbing many of these genes would manifest in *C. elegans* phenotypic profiles, we developed software to measure a comprehensive range of parameters while animals were subjected to an automated short-term habituation learning behavioral paradigm (**Figure 1E-I**, and see **Methods**). We chose to measure habituation and initial sensitivity to stimulation because habituation learning is impaired in individuals with ASD, and abnormalities in tactile sensitivity are present in >95% of cases.^2,69–77^ The degree of habituation impairment in ASD patients also correlates with the severity of social impairment, and recent studies in monogenic mouse models of ASD suggest peripheral tactile hypersensitivity and impaired habituation precede, and may even lead, to more complex cognitive and social impairments.^70,78^

To analyze the data we wrote custom scripts to extract 26 quantitative phenotypic features that fall into 5 categories of morphology, baseline locomotion, initial sensitivity, habituation learning, and spontaneous recovery/short-term memory retention (**Figure 1J** and see **Methods** and **Table S2** for a complete description of phenotypic features). The features we measured were designed to minimize redundancy while maintaining interpretability and keeping known biological information in mind. For example, we quantified multiple features of habituation as we have previously shown that they habituate to different extents and with different time courses (**Figure 1F-I**) and because there is growing evidence that different components of a single habituating response are mediated by different molecular mechanisms.^61,79,80^ Systematic quantification of the pairwise correlation between all possible phenotypic feature pairs revealed expected moderate correlations (e.g. between length and width) and clustering that reflected the feature categories we predefined (**Figure 1K**). The digital representations of all strains are freely available in their raw and processed forms (https://doi.org/10.5683/SP2/FJWIL8), allowing the code to be modified in subsequent analyses to extract any simple or compound phenotypic feature of choice.

### Quantitative phenotypic profiles identify shared functions among ASD-associated genes

There are many ways to visualize this data, but the most conceptually simple way is as a series of 26 quantitative reverse genetic screens, where each strain is compared to wild-type on each phenotypic feature to determine if and how that gene plays a role in the biological process of interest (**Figure 2 and online supplemental figure collection 1**). Phenotypes shared by a large number of ASD risk genes would be of particular interest, as they would hint at common biological functions among these seemingly diverse genes. We found that mutations in the vast majority of ASD-associated gene orthologs decrease multiple measures of size, most notably length (52/87, 60%), in age-synchronized populations (**Figure 2A**). Thus, the majority of genes putatively associated with ASD/neurodevelopmental disorders delay development or growth when inactivated.^81^

**Figure 2.**
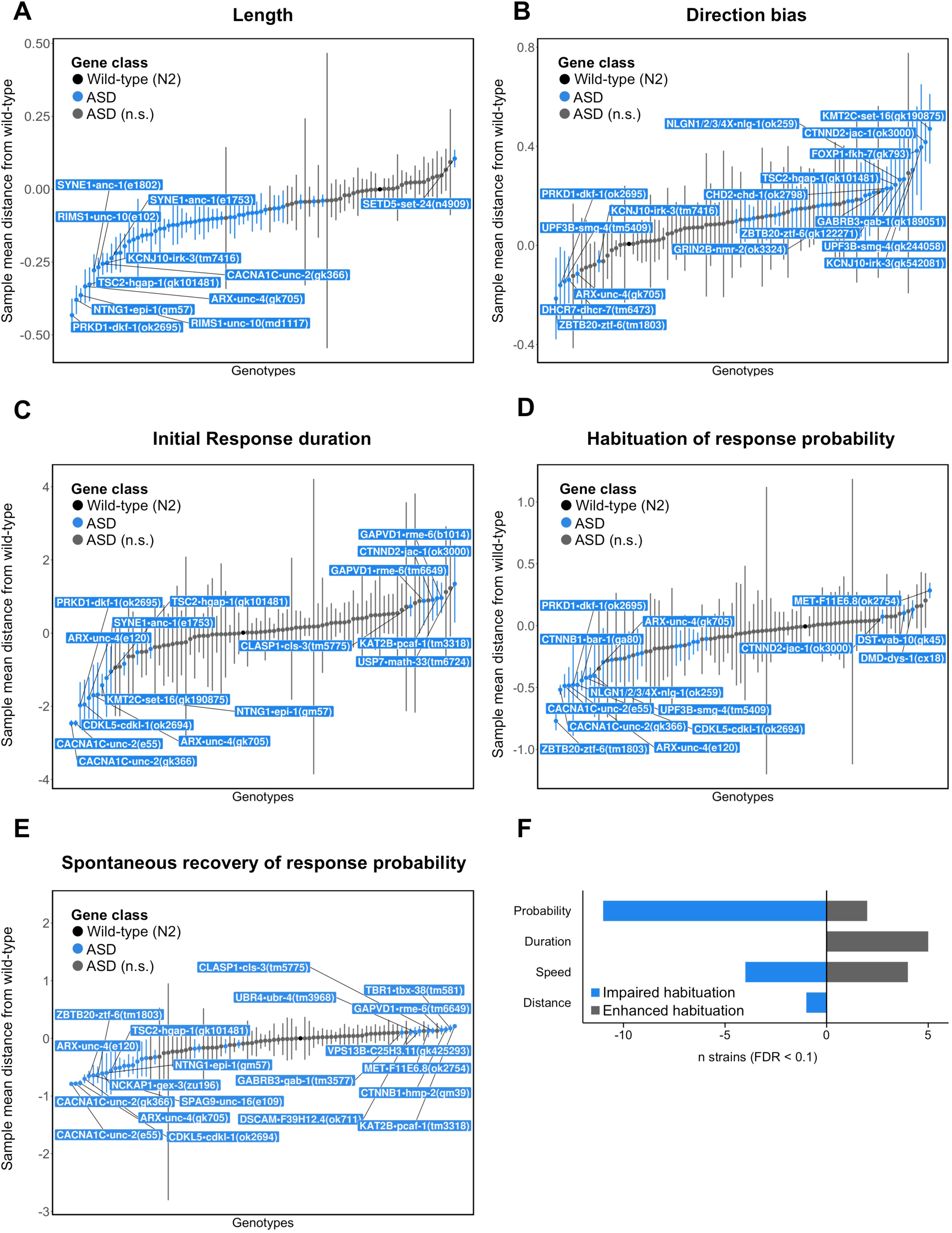
Quantitative phenotypic profiles enable rapid reverse genetic screens to identify shared functions among ASD-associated genes. All plots illustrate the sample mean distance of each genotype group from wild-type. Strains outside the 95% confidence interval of the wild-type distribution are labeled and colored blue. Only a maximum of ten strains are labeled in each direction per feature to prevent over plotting. **A)** Sample mean distance of length by genotype. The majority of ASD-associated gene orthologs decrease length when mutated. **B)** Movement direction bias by genotype. A large proportion of ASD-associated gene orthologs increase forward movement bias. **C)** Initial reversal response duration by genotype. ASD-associated gene orthologs are roughly equally likely to increase or decrease initial sensitivity to mechanosensory stimuli. **D)** Habituation of response probability by genotype. Many ASD-associated genes impair habituation of response probability **E)** Spontaneous recovery of response duration by genotype. Distinct partially overlapping sets of ASD-associated gene orthologs alter initial sensitivity, habituation learning, and spontaneous recovery/memory retention, indicating genetically dissociable underlying mechanisms. **A-E)** Error bars represent 95% confidence intervals. **F)** The number of strains with normal initial responses that either impair habituation (blue) or enhance habituation (gray) across the four habituating response metrics quantified.

Analysis of baseline locomotion features revealed that mutations in a large proportion of ASD-associated gene orthologs increase forward movement bias (32/87, 37%) and path length (distance travelled) compared to wild-type animals (**Figure 2B**). *C. elegans* normally spend the majority of their time in a forward locomotion behavioral state that is intermittently interrupted by brief spontaneous reversals.^82,83^ These two phenotypic changes together mean many strains with mutations in ASD-associated gene orthologs spend more time moving forward before eliciting a reversal. The frequency and duration of spontaneous reversals is modulated by the integration of multiple cross-modal sensory inputs,^82,84–93^ suggesting a widespread imbalance in the neural circuits that control spontaneous forward movement behavior toward increased activity in strains harboring mutations in orthologs of ASD-associated genes.

Assessing the duration, distance, and speed of reversal responses to the first mechanosensory stimulus indicated that mutations in orthologs of ASD-associated genes are approximately equally likely to result in initial hyper- or hypo-responsivity to tactile stimuli (**Figure 2C**; Note that bidirectional analysis of initial sensitivity was precluded only for response probability as there is a ceiling effect due to >90% of animals responding to the initial stimulus). These results identify novel positive and negative roles for multiple ASD risk genes in modulating mechanosensory processing.

Analysis of habituation learning across 87 strains revealed that mutations in many ASD-associated gene orthologs specifically impair habituation of response probability (**Figure 2D**). Indeed, even after filtering out strains with abnormal initial response probability ASD-associated gene orthologs were >5 fold more likely to impair habituation learning than enhance it (**Figure 2F and online supplemental figure collection 1**). This is a remarkably specific phenotype that causes the neural circuit to continue responding in an inflexible manner, as opposed to merely impairing the ability to detect stimuli or respond. Importantly, we only observed this phenomenon for response probability; there was no such consistent pattern of habituation impairment for the duration, distance, or speed of the same measured responses (**Figure 2F and online supplemental figure collection 1**). Taken together, these results suggest that many ASD-associated genes normally mediate plasticity of the likelihood, but not vigor, of responding to mechanosensory stimuli.

Finally, we discovered that distinct sets of genes alter initial sensitivity, habituation learning, and retention of the same component of the same behavioral response (**Figure 2C-E and online supplemental figure collection 1**). Adding to the complexity, we also discovered that different genes affect different components of the same behavior (e.g. some genes affect only response duration but not probability and vice versa). These results provide direct support for the hypothesis that habituation learning is controlled by several dissociable genetic mechanisms,^94,95^ and underscore the need to assess multiple complex phenotypes to gain a comprehensive understanding of the functions of ASD-associated genes.

### Phenotypic profiles of strains with mutations in ASD-associated genes define shared and unique functions and phenotypic modularity

As an alternative to visualizing the scores of all strains on each phenotype, one can instead visualize the scores of each strain on all phenotypes, or the ‘phenotypic profile’ for each strain (**Figure 3 and online supplemental figure collection 2**). Classical uncoordinated “unc” mutants, such as the calcium channel subunit *CACNA1C•unc-2(gk366)*, displayed the most severe overall quantitative phenotypic profiles, consistent with their documented roles in neuronal development and function (**Figure 3A**). Phenotypic profiles can also reveal previously unknown phenotypes even in well-characterized mutants. For example, β-catenin has well-known roles in development, so it follows that *CTNNB1•bar-1(ga80)* mutants are smaller in length and width than wild-type animals, but here we identify a previously unknown role for *CTNNB1•bar-1(ga80)* in habituation learning, evidenced by a profound deficit in habituation of response probability in *CTNNB1•bar-1(ga80)* mutants (**Figure 3B**). Animals carrying mutations in, *KAT2B•pcaf-1(tm3318)*, involved in CREB related transcription co-activation and acetyltransferase activity, have relatively normal baseline locomotion but profound alterations in sensory and learning behaviors that are only revealed through stimulation (**Figure 3C**). Conversely and more surprisingly, animals carrying mutations in genes such as *KCNJ10•irk-3(tm7416)*, an inward rectifying potassium channel, have relatively normal sensory and learning behaviors despite profoundly abnormal morphology and baseline locomotion (**Figure 3D**). Finally, there were many strains who were not significantly different from wild-type on almost all phenotypic features, such as the membrane palmitoylated protein encoded by *MPP6•C50F2.8(ok533)* (**Figure 3E**). Taken together these results indicate a remarkable degree of phenotypic modularity and provide a catalog of the unique phenotypic profiles of ASD-associated genes.

**Figure 3.**
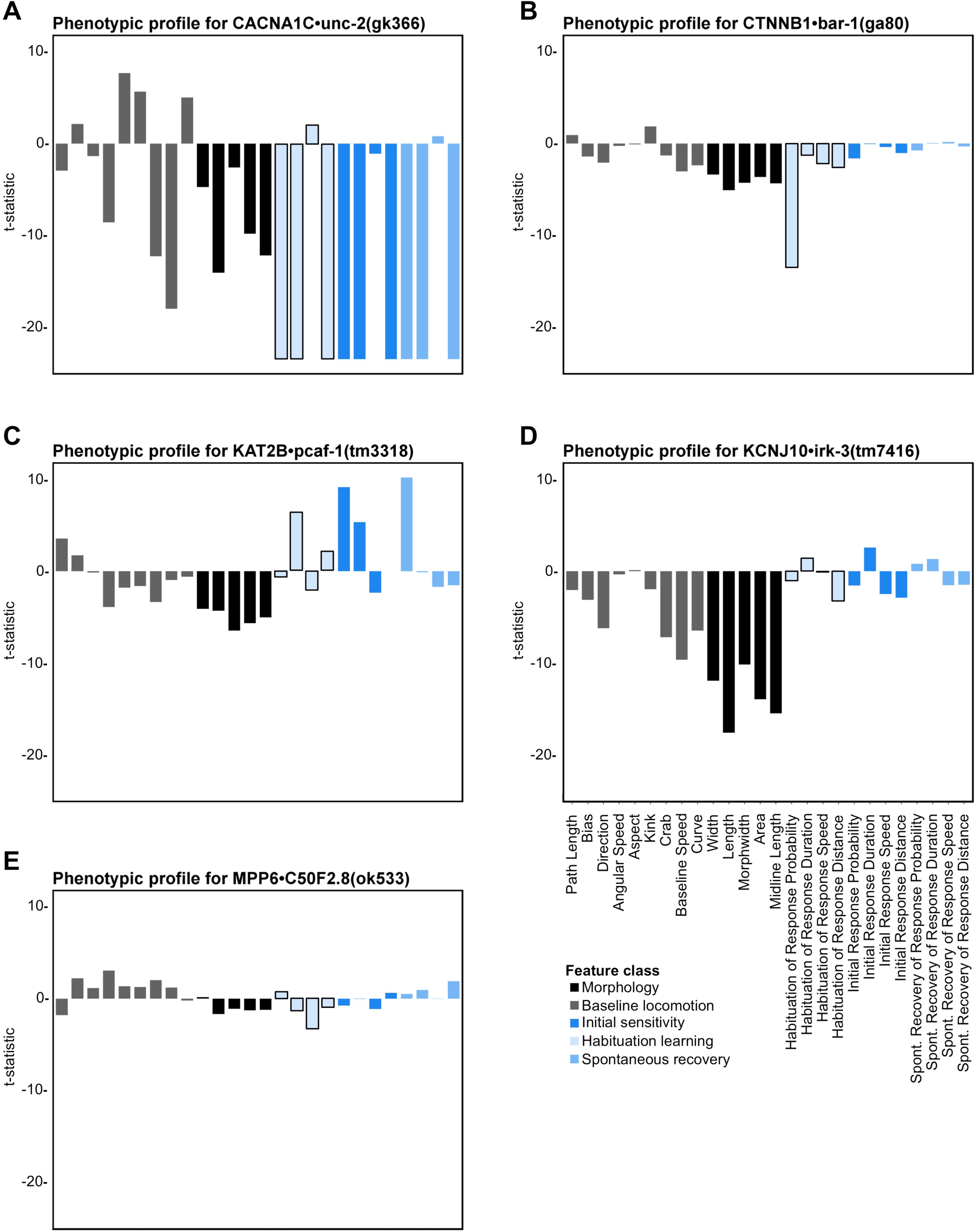
Phenotypic profiles of strains with mutations in ASD-associated genes define shared and unique functions and phenotypic modularity. For all plots bars represent directional t-statistics for each phenotypic feature listed across the x-axis in panel **D**. Color coding reflects feature classification. **A)** Phenoytpic profile for the classical uncoordinated mutant *CACNA1C•unc-2(gk366)* quantifies the extremity of phenotypic disruption across all logical phenotypic classes. **B)** Phenotypic profile for the heavily characterized *CTNNB1•bar-1(ga80)* strain confirms known roles in development (decreased length, width, etc.) and identifies a novel severe impairment in habituation of response probability. **C)** Phenotypic profile for *KAT2B•pcaf-1(tm3318)* exemplifying genes with relatively normal baseline locomotion behavior but more severe alteration in sensory and learning behaviors that could only be revealed through stimulation. **D)** Phenotypic profile for the inward rectifying potassium channel *KCNJ10•irk-3(tm7416)* exhibiting profound alterations in morphology and baseline locomotion yet relatively normal sensory and learning behaviors. **E)** Phenotypic profile for *MPP6•C50F2.8(ok533)* exhibiting relatively minor phenotypic alterations.

### A phenomic database of strains with mutations in ASD-associated genes

The most succinct and comprehensive way to visualize the data is as a phenomic heat map, illustrating the scores of all strains on all phenotypic features (**Figure 4A**). **Figure 4A** summarizes the scores of ∼18,000 animals (∼200 animals per genotype) across 87 genotypes and 26 phenotypes for a total of 2,262 *in vivo* genotype-to-phenotype assessments. These results show that for the phenotypes quantified and strains tested there is no single phenotype affected by all putative ASD-associated gene orthologs (**Figure 4A**). However, all strains were significantly different than wild-type on at least one phenotypic metric (**Figure 4B**). It is also clear that some phenotypes are more or less tolerant to mutation than others, for example a larger number of strains displayed altered morphological phenotypes compared to baseline locomotion, sensory or learning phenotypes (**Figure 4B**). Finally, our results indicate that distinct, partially overlapping sets of genes control the different classes of phenotypes (e.g. naïve sensitivity and habituation learning are influenced by different sets of ASD-associated genes) (**Figure 4A,B**).

**Figure 4.**
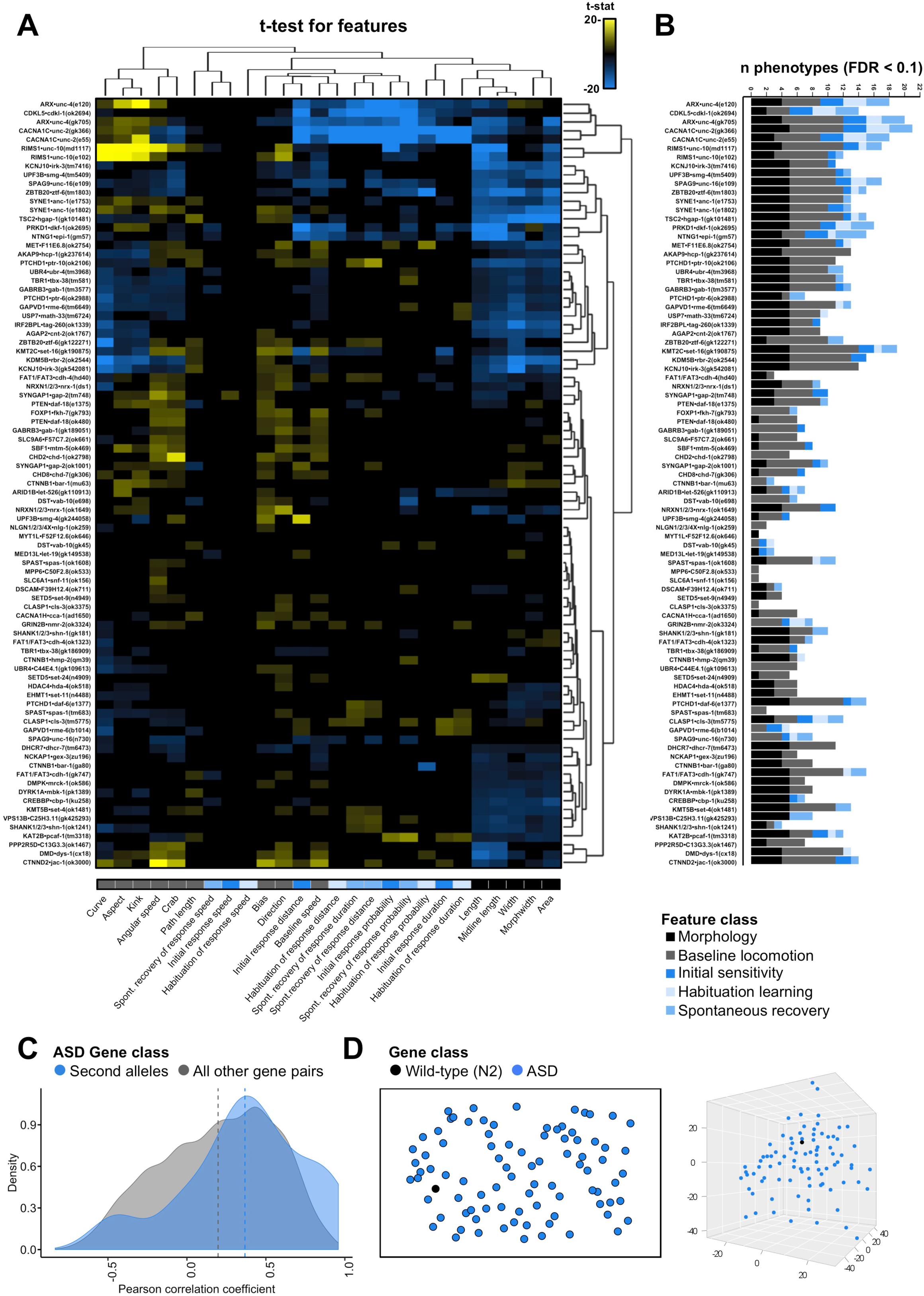
A phenomic database of strains with mutations in ASD-associated genes. **A)** Phenomic heat map summarizing the phenotypic profiles of 87 strains harbouring a mutation in an ortholog of an ASD-associated gene. Cells represent directional t-statistics from comparisons to wild-type controls. T-statistics are clipped at 20 and only cells significant at FDR < 0.1 are colored for ease of interpretation. A heat map illustrating the full range of all t-statistics for all strains can be found in **online supplemental figure collection 3**. **B)** The number of significantly different features for each strain tested. Stacked bars are color coded according to the five feature sub-classes. There was no single phenotype affected by every gene. All strains were significantly different from wild-type on at least one metric. **C)** Density plot illustrating the distribution of overall phenotypic profile Pearson correlation coefficients between second alleles in the same gene (21 pairs, blue) and all other possible gene pairs (3720 pairs, gray). **D)** 2D (left) and 3D (right) t-SNE plots illustrating the distance between the 87 strains harboring mutations in ASD-associated gene orthologs (blue) and wild-type (black) in phenotypic space.

Thus far, we have essentially conducted 2,262 exploratory analyses of which assayed phenotypes ASD-associated genes influence, and in doing so have identified hundreds of novel genotype-phenotype relationships. While there are many potential downstream uses for this database, we endeavored to illustrate a few of the most promising applications. First, we clustered the mutant strains based on their similarity in overall phenotypic profiles with the hypothesis, supported by recent large-scale model organism phenotyping efforts, that phenotypic similarity would enrich for functional genetic interactions among ASD-associated genes. However, while it has been successful for *C. elegans* morphology and baseline locomotion profiles in the past,^96–104^ a large-scale phenotypic clustering approach has not been attempted in combination with complex sensory and learning phenotypes. Prior to clustering we confirmed the sensitivity and consistency of our phenotypic measures by examining the correlation of pairs of alleles for genes we tested. Our sample contained second alleles for 21 genes. Analysis of the distribution of the overall phenomic correlations between second allele pairs revealed the average correlation was indeed higher than all other possible gene pairs (n = 3720) and that the second allele pair distribution was skewed towards highly positive correlations (**Figure 4C**).

With this assurance of specificity we then used several clustering methods to group genes based on phenotypic similarity to predict genetic interactions. Hierarchical clustering accurately identified several known interactions, such as those between the voltage-gated calcium channel *CACNA1C•unc-2* and the *Rab3* binding protein *RIMS1•unc-10* (**Figure 4A and online supplementary figure collections 3 & 4**).^105–109^ Our analysis also confirmed several recently discovered interactions, such as those between the dual specificity kinase *DYRK1A•mbk-1* and the histone acetyltransferase *CREBBP•cbp-1*, and predicted several novel interactions (**Figure 4A and online supplementary figure collections 3 & 4**).^110^ We then investigated the overall phenotypic architecture of ASD-associated genes. t-distributed Stochastic Neighbor Embedding (t-SNE) and multiple other clustering methods revealed that, while all strains were significantly different from wild-type, ASD-associated genes were largely continuously distributed in phenotypic space (**Figure 4D**). Indeed, there was no evidence for a small number of highly discrete clusters such that one could make a claim for distinct “phenotypic classes” of ASD-associated genes (**Figure 4A,D**).

We then used epistasis analysis to test some of our interactions predicted by phenotypic proximity *in vivo*. Using sensory and habituation learning features for hierarchical clustering revealed two high-confidence clusters with members that display impaired habituation of response probability as well as hyper-responsivity to mechanosensory stimuli (increased initial reversal response duration) (**Figure 5A-D and online supplementary figure collection 4**). Genes within these clusters were selected for epistasis analysis based on confidence of ASD-association and confirmation of genotype-to-phenotype relationships via analysis of a second allele or transformation rescue. Crossing between members of the same cluster revealed a novel functional interaction between the Chromodomain Helicase DNA Binding Protein *CHD8•chd-7(gk306)* and the GTPase-activating protein *GAPVD1•rme-6(b1014)*; the impairment in habituation of response probability of double mutants was not significantly different from single mutants, suggesting they function in the same genetic pathway to mediate short-term behavioral plasticity (**Figure 5E**). A second allele of *GAPVD1•rme-6(tm6649)* fell in the same cluster and displayed the same phenotypic profile as the crossed allele, and an additional allele of *CHD8•chd-7(gk209)*, tested after the initial large-scale characterization, also displayed the same phenotypic profile, confirming genotype-to-phenotype relationships (**Figure S5**). Importantly, *CHD8* is a high-confidence ASD-associated gene whereas *GAPVD1* is relatively low confidence, yet when they are inactivated in a model organism they cause strikingly similar phenotypic profiles and function together to promote normal habituation learning.

**Figure 5.**
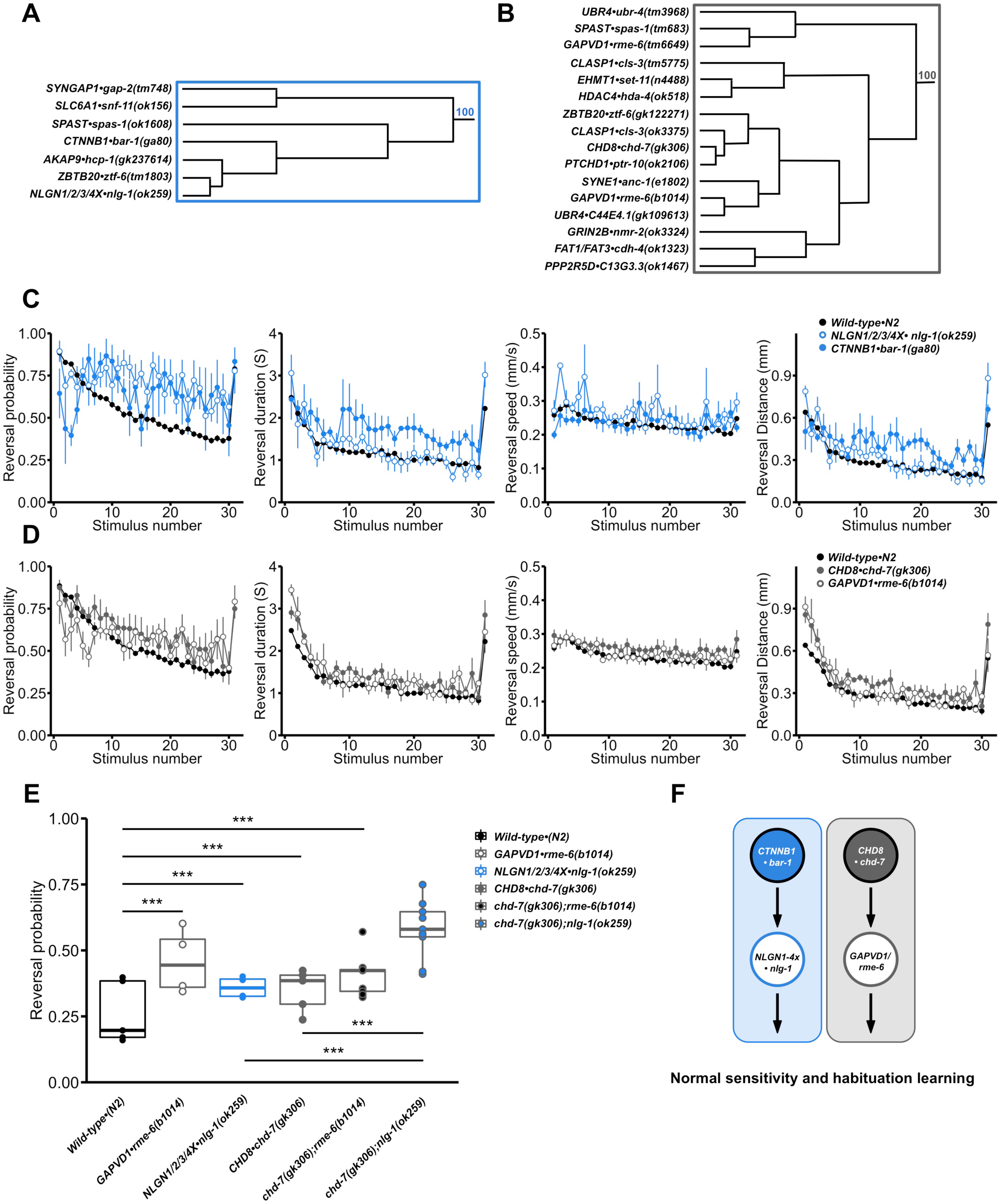
Combining phenotypic clustering and epistasis to map parallel genetic networks underlying hyper-responsivity and impaired habituation. **A&B)** Hierarchical clustering based on sensory and habituation learning features identified two sets of genes with members who display impairments in habituation of response probability and hyper-responsivity to mechanosensory stimuli (increased initial reversal response duration). Rectangles outline the largest clusters with AU p-values larger than 95%.^172^ **C)** Sensory and learning phenotypic profile for *NLGN1/2/3/4X•nlg-1(ok259)* and *CTNNB1•bar-1(ga80)* mutants. Each point represents a reversal response. Data are shown as mean±s.e.m using the number of plates as n. **D)** Sensory and learning phenotypic profile for *CHD8•chd-7(gk306)* and *GAPVD1•rme-6(b1014)* mutants. Each point represents a reversal response. Data are shown as mean±s.e.m using the number of plates as n. **E)** Final reversal probability across genotypes (average of the 28th-30th reversal response). *chd-7(gk306);rme-6(b1014)* double mutants do not display additive impairments in habituation, suggesting they function in the same pathway. *chd-7(gk306);nlg-1(ok259)* double mutants display additive effects on habituation of response probability suggesting they function in parallel pathways. Dots represent individual plate means, horizontal lines represent median of plate replicates. ***p<0.001, binomial logistic regression followed by Tukey’s HSD criterion was used to determine significance of the habituated level (proportion reversing at tap 30) for each pair of strains. **F)** Parallel genetic pathways of ASD-associated genes underlie mechanosensory hyper-responsivity and impaired habituation learning.

Crossing of mutations between clusters revealed that *CHD8•chd-7(gk306)* and the sole *C. elegans* ortholog of vertebrate neuroligins *NLGN1/2/3/4X•nlg-1(ok259)* function in parallel genetic pathways, with double mutants exhibiting additive impairments in habituation learning (**Figure 5E**). Interestingly, we also discovered a synthetic lethal interaction between *CHD8•chd-7(gk306)* and *CTNNB1•bar-1(ga80)*, suggesting that *CHD8* can function independently of its canonical role in mediating the Wnt/β-catenin signaling pathway.^111^ These results are consistent with the observation that Wnt/β-catenin targets are neither upregulated nor the cause of lethality in inviable *CHD8* homozygous knockout mice,^112^ suggesting Wnt independent functions of *CHD8* are conserved throughout evolution.^113^ We combined our results with the recent observation that Wnt/β-catenin signaling increases expression and synaptic clustering of *NLGN3*^114^ to draw parallel pathways underlying impaired habituation learning (**Figure 5F**). Taken together, these results demonstrate that systematic phenotypic clustering and epistasis analysis of complex sensory and learning phenotypes present a novel *in vivo* approach to map functional genetic network interactions among ASD-associated genes and prioritize candidates that would be missed by focusing on currently high-confidence genes.

### Phenomic profiles can be leveraged for *in vivo* variant functional assays

Another major challenge facing the genetics of ASD is the inability to interpret the functional consequences of the large number of rare variants of uncertain significance.^25,115–117^ Many of the variants found in ASD are so rare that they preclude traditional genetic approaches to infer pathogenicity.^34,118,119^ Indeed, even for genes for which we have a good understanding of their biological function it is often not possible to computationally predict the pathogenicity of a particular variant with the level of precision required in the clinic.^33,115,120,121^ Experimental variant functional assays, where the variant in question is introduced into a model system to observe its effects on function, can be combined with computational approaches to more clearly guide clinical assessment, but there is currently a paucity of such assays for most ASD-associated genes due to a lack of understanding of their normal biological functions.^118,122–126^ Here, we show how the phenomic functional data we generated by studying inactivating mutations in ASD-associated gene orthologs can be combined with the genetic tractability of *C. elegans* to develop transgenic rescue based *in vivo* variant functional assays using complex sensory and learning behaviors as a readout.

In our large-scale characterization we discovered that mutations in the sole *C. elegans* ortholog of vertebrate neuroligins cause impairments in habituation of response probability. We generated a wild-type *nlg-1::YFP* fusion transgene and found that expression of this construct in *nlg-1(ok259)* deletion mutants was sufficient to restore normal habituation learning behavior (**Figure 6 A-C**). We then used this transgenic rescue as a platform to assess the functional consequences of mutations equivalent to all conserved ASD-associated neuroligin missense variants by testing their ability to rescue, revealing varied partial loss-of-function (**Figure 6 D-G**). These functional results align with the effects of variants on three additional behavioral functional assays: octanal avoidance chemotaxis, thermotaxis, and sensory integration (**Figure 6H-K**). Each of these functional assays involve distinct neural circuits suggesting a general partial loss-of-function mechanism due to ASD-associated neuroligin variants in whole organism behavior. All ASD-associated variants were expressed at similar levels and properly localized to synapses *in vivo*, likely ruling out a simple pathogenic mechanism of improper trafficking or severely reduced cellular abundance (**Figure 6B**). Importantly, all ASD-associated gene orthologs, including those with no prior functional annotation, were significantly different from wild-type animals on at least one phenotypic metric in our characterization (**Figure 4B**), suggesting this will be a broadly applicable approach to decipher variants of uncertain significance.

**Figure 6.**
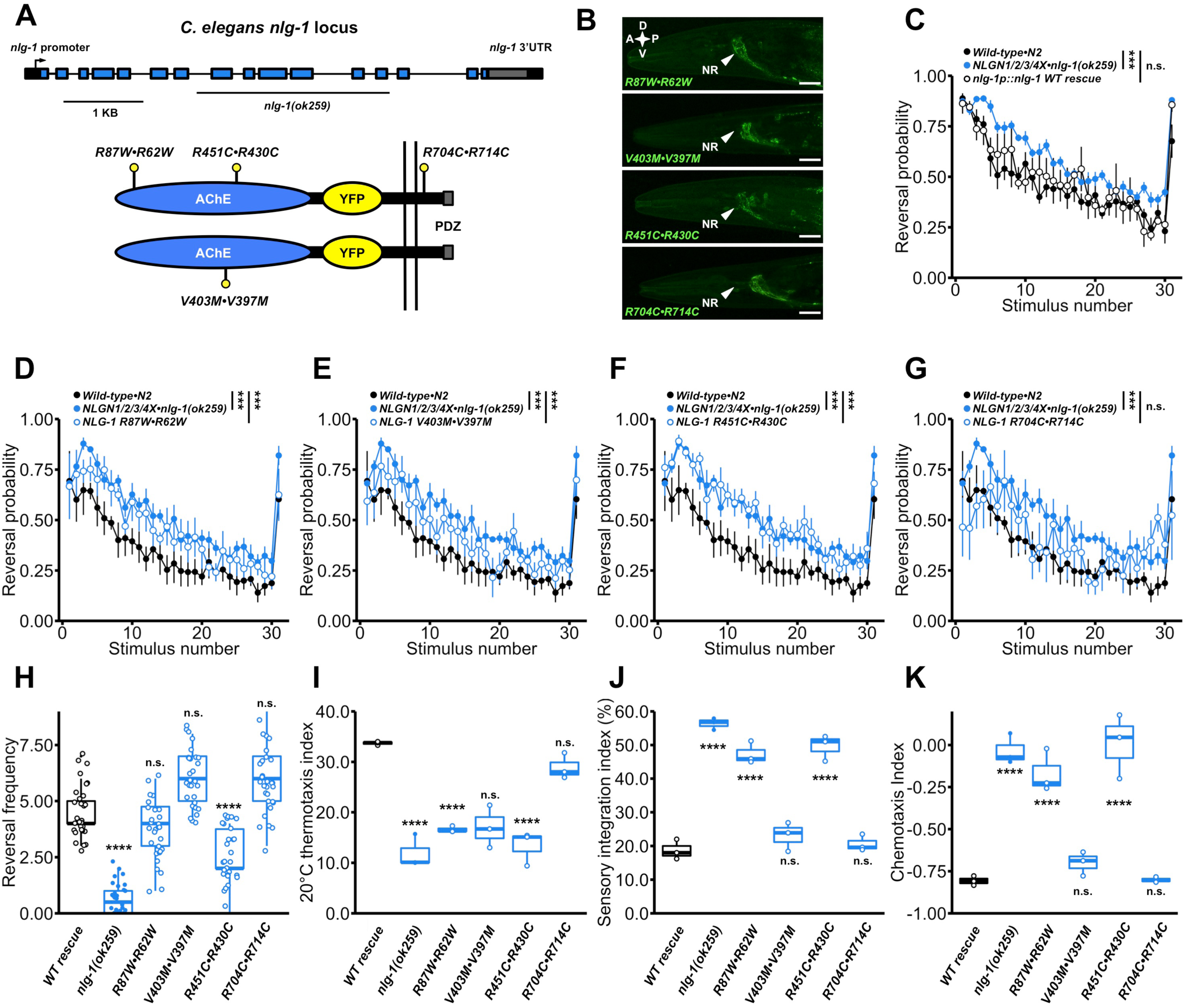
Functional assessment of ASD-associated missense variants in Neuroligins. **A)** Schematic illustration of the *nlg-1(ok259)* deletion allele and NLG-1::YFP fusion transgene used for transgenic rescue based variant functional assessments. Lollipops indicate the approximate locations of the equivalent ASD-associated missense variants assessed. Note that the R430C variant in *C. elegans* NLG-1 corresponds to a mutation in human *NLGN3* whereas the R62W, V397M, and R714C NLG-1 variants correspond to mutations in human *NLGN4*. **B)** All ASD-associated neuroligin variants were expressed at similar levels and localized to properly to synapses in the nerve ring and nerve cords. A = anterior, P = posterior, D = dorsal, V = ventral, NR = nerve ring, scale bar = 0.02 mm. **C)** Expression of wild-type NLG-1::YFP rescued *nlg-1(ok259)* deletion mutant impaired habituation of response probability. **D-G)** Each ASD-associated neuroligin variant was scored for its ability to rescue impaired habituation of response probability, revealing varied partial loss-of-function. **C-G)** Data are shown as mean±s.e.m using the number of plates as n. ***p<0.001, binomial logistic regression followed by Tukey’s HSD criterion was used to determine significance of the habituated level (proportion reversing at tap 30) for each pair of strains, n.s., not significant. **H-K)** Each ASD-associated neuroligin variant was scored for it’s ability to rescue spontaneous reversal frequency **(H)**, thermotaxis **(I)**, sensory integration **(J),** and octanal avoidance chemotaxis **(K)** in the *nlg-1(ok259)* deletion mutant. Dots represent individual worm reversal frequencies **(H)** or plate means **(I-K)**, horizontal lines represent median of plate replicates. **H-K)** ****p<0.0001, one-way ANOVA followed by Tukey’s HSD criterion was used to determine significance, n.s., not significant.

### CRISPR-Cas9 auxin inducible degradation reveals that *nlg-1* phenotypes are reversible by adult specific re-expression

Historically it was believed that the neurodevelopmental insults caused by monogenic risk factors for ASD were so severe that they would not be reversible in adulthood, and thus any treatment would have to be administered early to be effective.^127^ The seminal discoveries that phenotypes caused by mutations in *MECP2* could be reversed by transgenic expression of the protein in adulthood, after what had been presumed to be a critical developmental period had been missed, offered tremendous hope for families suffering from Rett syndrome.^128,129^ Conversely, the observation that inactivation of *MECP2*, Neuroligins, and several other ASD-associated genes in adulthood could cause severe electrophysiological and behavioral impairments demonstrated that they were not simply neurodevelopmental genes, and instead that they had ongoing functions throughout the lifespan to promote normal sensory and learning behaviors.^130–133^ This information must be taken into account for any therapy designed to treat these monogenic forms of ASD. These and other conditional rescue and inactivation experiments are valuable to ASD research, but they are expensive, technically difficult, and time consuming in mammalian model systems and are thus not yet practical for many ASD-associated genes. With *C. elegans*, however, conditional and reversible protein depletion is precise, rapid, and straightforward owing to their genetic tractability and the recent advent of CRISPR-Cas9 Auxin Inducible Degradation (AID).^134,135^ AID relies on transgenic expression of TIR1, an inducible E3 ubiquitin ligase that targets any protein with an AID degron peptide tag for degradation by the proteasome only when in the presence of its necessary activating hormone auxin (**Figure 7**). Moreover, upon removal from auxin TIR1 is inactivated, allowing rapid reconstitution of protein expression in large populations of freely behaving and intact animals without the need for surgery or vector delivery.^134^

**Figure 7.**
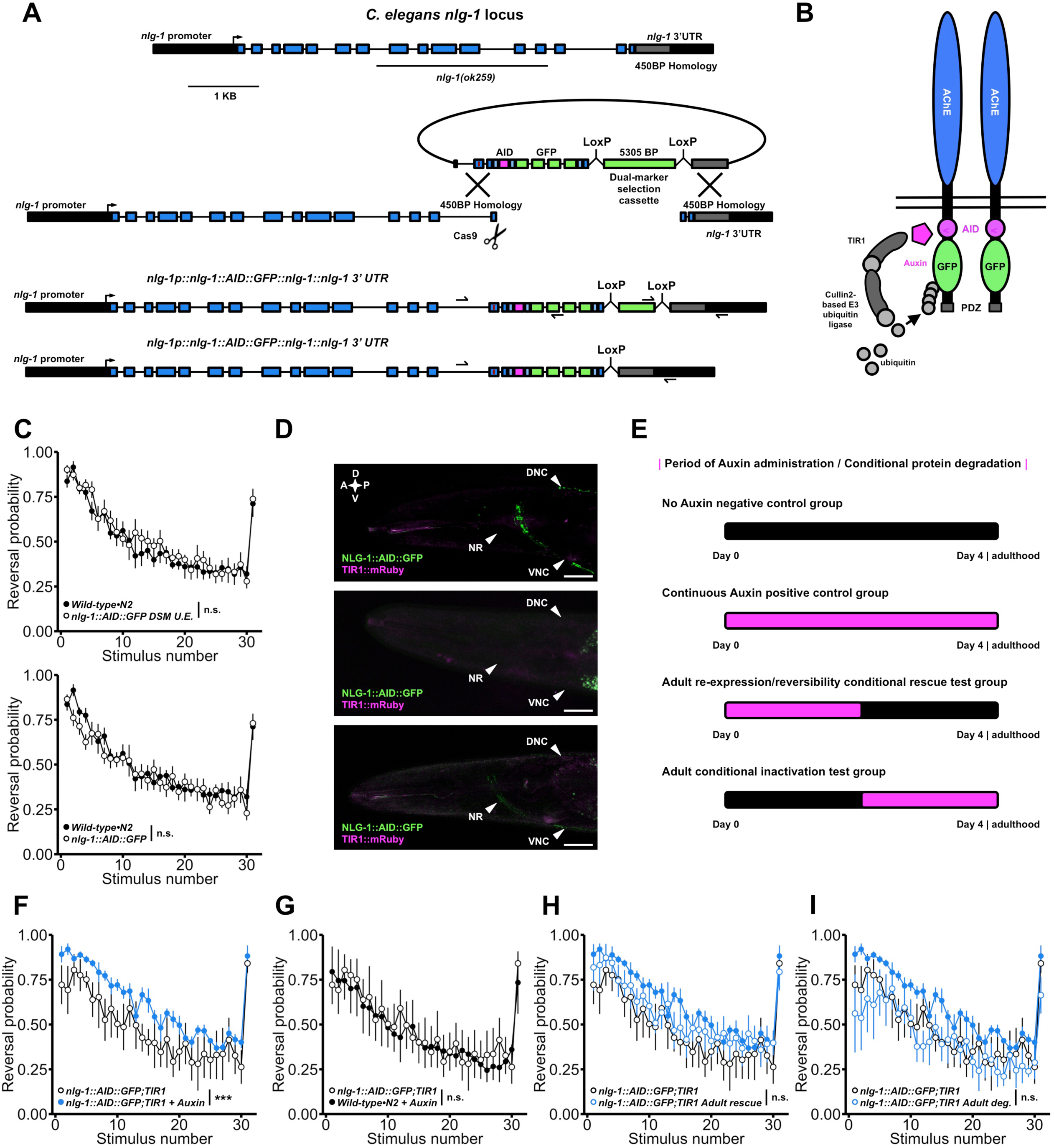
CRISPR-Cas9 auxin Inducible Degradation reveals phenotypes caused by developmental loss of neuroligin can be rescued by adult re-expression. **A)** Schematic illustration of the modified Dual Selection Marker (DSM) Cassette^54^ CRISPR-Cas9 genome editing strategy used to insert GFP and a short degron peptide tag into the endogenous neuroligin locus. The maroon line in the upstream homology arm of the repair template indicates the location of an engineered silent mutation in the protospacer adjacent motif to prevent cleavage of the exogenous DNA. **B)** Schematic illustration of the CRISPR-Cas9 engineered NLG-1::AID::GFP transgene used for AID experiments. In the presence of auxin TIR1 (a transgenically-expressed E3-ubuiqitin ligase) is activated targeting the fusion protein for degradation. Following auxin treatment TIR1 is inactivated allowing straight-forward conditional degradation and re-expression of the fusion protein. **C)** The fusion protein is fully functional; *NLG-1::AID::GFP* animals did not display habituation impairments before (top) or after (bottom) DSM selection cassette excision. **D)** The fusion protein localizes properly to synapses in the nerve ring and nerve cords (top). Treatment with 0.025 mM auxin is sufficient for complete degradation of the fusion protein (middle) that is reversible 48 hours after removal from auxin (bottom). A = anterior, P = posterior, D = dorsal, V = ventral, NR = nerve ring, scale bar = 0.02 mm. **E)** Schematic illustration of the time period of auxin administration for each of the conditional inactivation and rescue groups tested. **F)** Continuous auxin administration recapitulated induced impairments in habituation of response probability. G) *Wild-type•N2* animals continuously treated with auxin exhibited normal habituation. **H)** Adult specific re-expression of neuroligin partially rescued impaired habituation of response probability. **I)** Adult specific degradation of neuroligin did not induce habituation impairments. **C, F-I)** ***p<0.001, binomial logistic regression followed by Tukey’s HSD criterion was used to determine significance of the habituated level (proportion reversing at tap 30) for each pair of strains, n.s., not significant. Data are shown as mean±s.e.m using the number of plates as n.

Despite extensive study, transgenic rescue tests of phenotypic reversibility have not been completed for any member of the vertebrate neuroligin family.^107,136,137^ We used the CRISPR-Cas9 Dual Selection Marker Cassette genome editing strategy^54^ to insert GFP and a short AID degron peptide tag into the endogenous *C. elegans* neuroligin locus 13 residues before the stop codon (**Figure 7A**). This fusion protein localized properly to synapses and is fully functional – there were no habituation impairments or gross neuroanatomical abnormalities in genome-edited animals (**Figure 7C,D**). We then crossed this strain into animals expressing TIR1 under a ubiquitous promoter and observed that treatment with 0.025mM auxin for <10.0 hours was sufficient for complete degradation of the functional NLG-1::AID::GFP fusion protein at multiple life stages (**Figure 7D**). Moreover NLG-1::AID::GFP was partially recovered 48 hours after removal from auxin (**Figure 7D**). We used this approach to test several conditional rescue and inactivation groups simultaneously (**Figure 7E**). Animals reared on auxin for continuous degradation displayed habituation impairments equivalent to those of *nlg-1(ok259)* null mutants, confirming effective degradation (**Figure 7F**). Importantly, wild-type animals continuously exposed to auxin displayed no morphological or behavioral abnormalities (**Figure 7G**). Strikingly, we observed that for animals reared on auxin to degrade NLG-1 throughout development, adult specific expression of neuroligin was sufficient to partially rescue the habituation impairment phenotype (**Figure 7H**). Surprisingly, adult-specific degradation of neuroligin did not lead to impaired habituation learning behavior (**Figure 7I**). These results indicate a critical role for neuroligin in generating a circuit properly tuned for normal mechanosensory processing and short-term behavioral plasticity. Further, they suggest that behavioral disruptions caused by developmental loss of neuroligins might be at least partially reversible by adult expression. Given the speed of machine vision phenotyping, and relative ease of CRISPR-Cas9 genome editing in *C. elegans,* AID represents a scalable approach that can be applied to diverse ASD-associated genes.

## DISCUSSION

In total we quantified 26 phenotypes for >18,000 animals across 87 genotypes and identified hundreds of novel genotype-phenotype relationships. Our database approximately doubles the number of animals in a recent aggregation of all other machine vision behavioral datasets collected in this organism to date,^138^ and provides the first such systematic analysis that includes both complex sensory and learning phenotypes. By making the raw and processed data available online, we have created open and shareable phenotypic atlas of *C. elegans* strains carrying mutations in orthologs of ASD-associated genes. We have shown how this database can be used to identify shared functions, map genetic networks, gauge missense variant effect, and determine reversibility of phenotypic disruptions.

We found that the vast majority of strains with mutations in ASD-associated genes displayed delayed development, a finding consistent with a recent large-scale analysis of inactivating mutations in constrained genes in humans.^81^ As the number of genes assessed increases it will be interesting to conduct a pathway enrichment analysis to determine if mutations in particular pathways lead to ASD relevant phenotypes with or without developmental delay or whether particular phenotypes are more characteristic of genes predominantly associated with ASD versus Intellectual Disability/Developmental Delay (ID/DD).^25,101^ We also found that a large number of ASD-associated genes impair habituation, a finding that is consistent with a recently reported large-scale analysis of ASD- and ID/DD-associated genes in *Drosophila* and humans.^67^ Interestingly, our data show for the first time that ASD-associated genes specifically impair habituation of response probability. These results provide a genotype-to-phenotype list (**Figure 2D**) that may hint at a common pathological mechanism that impairs plasticity of a neural circuits’ decision to respond without altering response vigor. Indeed, these *Drosophila* and *C. elegans* results likely reflect a behavioral outcome of circuit-level hyperexcitability recently discovered in several human iPSC derived neuronal culture models of a number of monogenic ASD risk factors.^68,139^ The results also provide a plausible explanation for inconsistent reports of impaired habituation in humans, which variably employ diverse response metrics most often without genetic stratification of patient populations.^69^ While these shared phenotypes are exciting, our results also reveal a remarkable diversity and modularity in phenotypic disruptions, and thus clearly indicate that single phenotype *in vivo* functional validation and characterization efforts will be insufficient to capture the complex multi-facetted phenotypic disruptions that stem from inactivating mutations in ASD-associated genes.

### Phenomic clustering and epistasis to map genetic networks among ASD-associated genes *in vivo*

We combined clustering based on similarity in phenotypic profiles with epistasis analysis to map functional genetic networks among ASD-associated genes and identified novel parallel genetic networks centered on *CHD8•chd-7* and *NLGN1/2/3/4X•nlg-1* that underlie hyper-responsivity and impaired habituation learning. These results provide *in vivo* functional support to the proposed broad categorization of ASD-associated genes into those involved in synaptic neuronal communication (neuroligin) and gene expression regulation (*CHD8*).^15,25,140^ An exciting question for future research will be to determine how well the phenomic functional interactions delineated here map on to biochemical interactions. Indeed, *in vivo* genetic networks will serve as an important benchmark from which to compare and extend existing networks of ASD-associated genes. Phenomic clustering will be particularly useful for capturing long-range functional interactions between proteins expressed in different cells or even different points in development, which cannot be detected by measures of direct protein-protein interactions or co-expression.

### ASD-associated genes are continuously distributed in phenotypic space

We found that the set of strains we characterized were continuously distributed in phenotypic space, and that there were no clear highly separated discrete phenotypic clusters. Even in this rigorously controlled genetic background and environment there is no basis for a classification of ASD-associated genes into distinct ‘phenotypic classes.’ This would suggest that the inability to cluster the heterogeneous continuously distributed clinical population based on shared phenotypes or symptoms does not necessarily stem from difference in genetic background or environmental exposures. This is in contrast to the molecular level, where multiple studies and our present work suggest there are functionally distinct classes of ASD-associated genes.^3,25,31^ While phenotypic profiles were continuously distributed, certain genes were closer to each other than others in a manner that reflected underlying molecular interactions. The observation that ASDs are a group of etiologically distinct and variably phenotypically similar disorders provides further motivation for tailored treatments designed on the bases of shared molecular pathway disruptions.^3,31^

It is also important to consider whether the functions of ASD-associated genes discovered here will be conserved to their human orthologs, and even if they are conserved, whether they display any specificity to ASD. Our data and the literature show that there is no single phenotype at any level that can definitively classify an ASD-associated gene from those causing related disorders. Regardless of the qualitative diagnostic label applied, understanding the function of these poorly characterized neurodevelopmental disorder-associated genes will be extremely valuable for understanding pathology and designing treatments. Several of the shared functions for ASD-associated genes identified here, such as promoting normal development or habituation of response probability, have concurrently been discovered in higher model organisms and human iPSCs, suggesting that many of the other novel functions reported here will also be conserved and relevant to ASD. It is also worth reiterating that many human genes have repeatedly been shown to be able to functionally replace their *C. elegans* orthologs. For example, human *NLGN1* and *NLGN4* have both been shown to rescue multiple sensory abnormalities caused by loss of *C. elegans nlg-1*.^49,141^ We have also recently shown that directly replacing *daf-18* with only a single copy of its human ASD-associated ortholog *PTEN* at the endogenous locus using CRISPR-Cas9 is able to rescue multiple sensory abnormalities caused by complete deletion of *daf-18*.^142^ A recent systematic humanization effort of 414 genes in yeast showed that >50% of human genes could functionally replace their yeast ortholog.^143^ Indeed, at this point it would not be surprising if a highly conserved human gene could replace its *C. elegans* ortholog. All model systems have their relative strengths and weaknesses, and the fastest and most generalizable insights in ASD research will undoubtedly come from synthesis of large amounts information derived from diverse model systems and patients.

### Phenomic profiles can be leveraged for *in vivo* assays of missense variant effect

We used neuroligin as a proof-of-principle to show how our phenomic profiles can be leveraged to establish *in vivo* variant functional assays. We found that all variants tested displayed varied partial loss-of-function despite effective expression and proper subcellular localization. Further, we observed that variant functional results were consistent across multiple behavioral functional assays involving distinct neural circuits. While the mechanisms underlying the partial loss-of-function behavioral effects of ASD-associated neuroligin variants currently remains controversial,^144–149^ these results provide clinically relevant knowledge, as a rationally designed treatment should be designed to increase neuroligin function while taking into account the presence of an existing dysfunctional protein. Importantly, all strains were significantly different from wild-type on at least one metric, opening the door for many diverse transgenic rescue-based *in vivo* variant functional assessments moving forward. Many genes were significantly different on multiple metrics, giving researchers with interests in particular biological processes the ability to choose a phenotypic functional assay most suited to their needs. Our results provide a critical first step in establishing variant functional assays by systematically characterizing genotype-phenotype relationships that can be confirmed by transformation rescue or CRISPR-Cas9 knockout. A strength of *C. elegans* will continue to be the ability to rapidly assess multiple variants in complex sensory and learning behaviors *in vivo*.

### Harnessing phenotypic profiles for tests of adult reversibility

We found adult expression of neuroligin could partially reverse the hyper-responsivity and impaired habituation phenotypes of animals that developed without neuroligin. Interestingly, we also found adult specific inactivation did not produce phenotypic disruptions. These results are surprising, as inactivation of *NLGN1*, *NLGN2* and *NLGN3* in mature vertebrates produce abnormalities in several complex behaviors.^130,131,150^ However, a plausible explanation could be that neuroligin is only necessary in adulthood for forms of learning and memory that require structural plasticity. Indeed, all studies reporting an adult requirement for neuroligin examined different forms of long-term memory, which in contrast to the short-term memories studied here, require *de novo* activity-dependent synapse growth and maturation. Further, the phenotypes observed often did not manifest until weeks after neuroligins were inactivated in adult animals.^151^ These results lead to a model in which neuroligin is sufficient to build a circuit capable of normal sensitivity and short-term habituation at any point throughout the lifespan, but once that circuit has been built its function is no longer required for short-term learning. Continued function of neuroligin would then remain necessary only for more complex forms of long-term learning and memory that require *de novo* activity-dependent growth and maturation of synaptic connections. There is no evidence yet of new synapse formation underlying long-term learning in *C. elegans,* making this model currently difficult to test. However, an adult form of experience/activity-dependent neural circuit remodeling has just been discovered in *C. elegans*, and neuroligin was identified a critical component in this process.^152^ More importantly for individuals with ASD, these results suggest that some phenotypic alterations due to developmental loss of neuroligin may be reversible in adulthood, and provide a rapid and inexpensive strategy where AID can be used to test reversibility of phenotypic disruptions caused by diverse ASD-associated genes.

### Conclusions

We have provided the first systematic *in vivo* phenomics analysis of ASD-associated genes and identified shared delays in development, hyperactivity, and impairments in habituation learning. Our data adds to the recent and rapidly expanding use of model organism phenomics to discover the functions of poorly characterized genes identified through genomic sequencing.^44,101,153–158^ We have shown how such data can be used to identify novel genetic interactions, establish variant functional assays, and develop tests of phenotypic reversibility. There is substantial evidence that an insufficient understanding of the biology of many disease-associated genes has prevented the successful development of therapies and that preclinical research is biased towards experimentally well-accessible genes.^159–163^ It is ideal to be systematic and unbiased whenever possible, an opportunity high-throughput model organisms such as *C. elegans* afford and thus one we should continue to exploit. As we continue to chart the phenotypic landscape of ASD-associated genes, the complicated paths to understanding mechanisms and developing personalized treatments become simpler to navigate.

## STAR Methods

### Key Resource Table (separate document)

#### Contact for Reagent and Resource Sharing

Further information and requests for resources and reagents should be directed to and will be fulfilled by the Lead Contact, Catharine H. Rankin.

#### Ortholog identification and strain selection

*C. elegans* orthologs of human ASD-associated genes were identified by querying OrthoList using Ensembl gene IDs, as previously described.^45^ **Table S1** Describes all Orthology relationships used in this study. During the course of this project The Alliance of Genome Resources (https://www.alliancegenome.org/) created a web tool that allows for identification of the “best” matched Ortholog, defined as the ortholog predicted in the queried species by the highest number of gold-standard algorithms.^47^ The vast majority of the *C. elegans* orthologs predicted by OrthoList (85%) are also defined as the best ortholog by this new tool (**Table S1**). OrthoList was also updated to OrthoList2 during the course of this work. Again there is large agreement between our orthology predictions and those made using the recently updated OrthoList2 (95% of OrthoList relationships are supported by OrthoList2; **Table S1**). In situations where multiple human ASD-associated genes share a single *C. elegans* ortholog predicted by OrthoList the single *C. elegans* gene was used to study the larger vertebrate family, as has been done previously.^137,164,165^ For example, *nlg-1* and *shn-1* are the sole *C. elegans* orthologs of all vertebrate neuroligin and shank family proteins, respectively (**Table S1**). Note that throughout the manuscript the “*•*” symbol is used to represent the human-to-*C. elegans* orthology relationship of interest, e.g. *GAPVD1•rme-6(b1014)*.

Strains harboring mutations in orthologs of ASD-associated genes were identified using WormBase and ordered from the *Caenorhabditis* Genetics Centre (CGC), the National BioResource Project of Japan (NBRP), or received from a collaborator following a formal request. Strains carrying putative null and loss-of-function alleles were prioritized. In many cases multiple strains harboring distinct loss-of-function alleles were characterized. Where such alleles were not available, or where null alleles are known to result in lethality or fecundity defects severe enough to impede high-throughput characterization in our assay a strain carrying a known or predicted deleterious missense mutation was characterized instead. The complete list of all strains and alleles used in the work are described in **Table S1**.

#### Animals

Strains were maintained on NGM (nematode growth medium) plates seeded with E. coli strain OP50 according to standard experimental procedures. 96h post-hatch hermaphrodite animals were used for all experiments. A complete list of strains used is available in **Table S1** and the **Key Resources Table**.

#### Microbe strains

The Escherichia coli OP50 strain was used as a food source for C. elegans.

#### Behavioral assays

For the mechanosensory habituation paradigm animals were cultured on Nematode Growth Medium (NGM) seeded with *Escherichia coli* (OP50) and age synchronized for behavioral tracking as described previously.^79,166,167^ Animals were synchronized for behavioral testing on Petri plates containing Nematode Growth Media (NGM) seeded with 50 µl of OP50 liquid culture 12-24 hours before use. Five gravid adults were picked to plates and allowed to lay eggs for 3-4 hours before removal. For all Multi-Worm Tracker experiments 3-6 plates (20-100 worms/plate) were run for each strain on each testing day. The animals were maintained in a 20°C incubator for 96 hours.

Our behavioral paradigm (**Figure 1E**) consisted of a 5-minute period to recover from being placed on the tracker followed by a 5 min baseline period from which we computed multiple measures of morphology and baseline locomotion. Beginning at 10 minutes we administered 30 mechanosensory stimuli to the Petri plate holding the animals at a 10 second interstimulus interval (ISI) using an automated push solenoid (**Figure 1A and E**). Animals respond to a mechanosensory stimulus by emitting a reversal response (crawling backwards) allowing us to assess multiple measures of naïve sensitivity (e.g. reversal likelihood, duration, etc.) (**Figure 1E-I**). With repeated stimulation there is a decrease in the likelihood of a reversal, as well as the duration, speed, and distance of reversals (habituation). Following habituation training, we allowed a 5 minute recovery period after which we administered a 31^st^ stimulus to gauge spontaneous recovery from short-term habituation - an assay of short-term memory retention (**Fig 1E-I**).

Standard sensory integration, octanal avoidance, and thermotaxis assays were conducted as previously described.^141,168^

#### Multi-Worm Tracker behavioral analysis and statistics

Multi-Worm Tracker software (version 1.2.0.2) was used for stimulus delivery and image acquisition.^60^ Phenotypic quantification with Choreography software (version 1.3.0_r103552) used “--shadowless”, “--minimum-move-body 2”, and “--minimum-time 20” filters to restrict the analysis to animals that moved at least 2 body lengths and were tracked for at least 20 s. Standard choreography output commands were used to output morphology and baseline locomotion features.^60^ A complete description of the morphology, baseline locomotion, sensory, and habituation learning features can be found in the Multi-Worm Tracker user guide (https://sourceforge.net/projects/mwt/)^60^ and **Table S2**. The MeasureReversal plugin was used to identify reversals occurring within 1 s (d*t* = 1) of the mechanosensory stimulus onset. Comparisons of “final response” comprised the average of the final three stimuli. Custom R scripts organized and summarized Choreography output files. No blinding was necessary because the Multi-Worm Tracker scores behavior objectively. For the initial large-scale characterization (**Figures 1-4**), features were pooled across plate replicates for each mutant strain and means were compared to the mean of the wild-type distribution with an unpaired t- test and Benjamani-Hochberg control of the false discovery rate at 0.1.^169^ Final figures were generated using the ggplot2 package in R.^170^ For targeted confirmation and follow-up analyses (**Figures 5-7**) responses were pooled across plates and compared across strains using binomial logistic regression for habituation or one-way ANOVA for additional behavioral assays (**Figure 6 H-K**) with Tukey’s honestly significant difference (HSD) criterion as previously described.^167,171^ Each targeted confirmation and follow up experiment was independently replicated at least twice. Alpha values of 0.001 or 0.0001 were used to determine significance for Logistic regression and one-way ANOVA statistical tests respectively. Final figures were generated using the ggplot2 package in R. For all Multi-Worm Tracker experiments 3-6 plates (20-100 worms/plate) were run for each strain on each testing day. Sample sizes for each behavioral assay were chosen to be either equal to or greater than sample sizes reported in the literature that were sufficient to detect biologically relevant differences.

All raw and processed data, and the results of all statistical tests can be found at (https://doi.org/10.5683/SP2/FJWIL8), all analysis code is freely available at (https://github.com/PavlidisLab/McDiarmid-etal-2019_Multi-Worm-Tracker-analysis).

#### Clustering analyses

The Student’s T-statistic was used as a numerical score to represent the difference between wild-type and mutant animals for each phenotypic feature; this created a numerical profile of phenotypic features for further analysis. All clustering analyses were performed in R. Correlation distributions were visualized using ggplot2. Average-linkage hierarchical clustering was performed with pvclust using correlation as the distance measure, and 50,000 rounds of bootstrapping.^172^ t-SNE clustering was performed using the Rtsne package^173^ with hyperparameters: initial principal component analysis = TRUE, perplexity = 10, and theta = 0 for the final 2D and 3D visualizations. The dendrogram and heat maps were visualized with pheatmap^174^ for the correlation matrix heat map, iheatmapr^175^ for the phenomic profile heat maps, and pvclust for the dendrograms.

### List of strains generated in this work

The following strains were created for this work either via standard genetic crosses for double mutants, or via microinjection of plasmid DNA for the generation of extrachromosomal array or CRISPR-Cas9 genome engineered transgenic lines:

VG870-871 *chd-7(gk306) I; nlg-1(ok259) X*

VG872-873 *chd-7(gk306) I; rme-6(b1014) X*

RM3540 *pha-1(e2123ts); nlg-1(ok259); mdEx1035[Pnlg-1 v2.1::NLG-(R62W)1::YFP v3; pBX]*

RM3516-17 *pha-1(e2123ts); nlg-1(ok259); mdEx1016-1017[Pnlg-1 v2.1::NLG-1(V397M)::YFP v3; pBX]*

RM3536 *pha-1(e2123ts); nlg-1(ok259); mdEx1033[Pnlg-1 v2.1::NLG-1(R430C)::YFP v3; pBX]*

RM3537 *pha-1(e2123ts); nlg-1(ok259); mdEx1034[Pnlg-1 v2.1::NLG-1(R714C)::YFP v3; pBX]*

VG880 *nlg-1(yv15[nlg-1p::nlg-1::AID::GFP:: + LoxP pmyo-2::GFP::unc-54 UTR prps-27::NeoR::unc-54 UTR LoxP + nlg-1 3’ UTR])*

VG881 *nlg-1(yv16[nlg-1p::nlg-1::AID::GFP:: + LoxP + nlg-1 3’ UTR])*

VG890-891 *nlg-1(yv16[nlg-1p::nlg-1::AID::GFP:: + LoxP + nlg-1 3’ UTR]); ieSi57[eft-3p::TIR1::mRuby::unc-54 3’ UTR + cbr-unc-119(+)] II; unc-119(ed3) III*

#### Strain and plasmid generation

The neuroligin missense variant plasmids were constructed by performing standard site-directed mutagenesis on our previously described *nlg-1* YFP functional fusion protein construct derived from the yk497a9 cDNA.^168^

The Moerman lab guide selection tool (http://genome.sfu.ca/crispr/) was used to identify the *nlg-1* targeting sgRNA.^53^ The *nlg-1* sgRNA sequence: TCACCAACGTGTCCACGTCA was cloned into the *pU6::klp-12* sgRNA vector (obtained from Calarco lab) using site-directed mutagenesis and used for all editing experiments. The *nlg-1::AID::GFP::nlg-1* upstream and *nlg-1 3’ UTR* downstream homology arms were synthesized by IDT and cloned into the loxP_myo2_neoR repair construct (obtained from Calarco lab) using Gibson Assembly.

*C. elegans* wild-type N2 strain was used for all CRISPR-Cas9 editing experiments. Genome edits were created as previously described.^54^ In brief, plasmids encoding sgRNA, Cas9 co-transformation markers pCFJ90 and pCFJ104 (Jorgensen lab, Addgene) and the selection cassette flanked by homology arms (∼500 bp) containing *AID::GFP* were injected into wild-type animals. Animals containing the desired insertions were identified by G418 resistance, loss of extrachromosomal array markers, and uniform dim fluorescence of the inserted GFP.

#### Genotype confirmation

Correct insertion of the *GFP::AID sequence* was confirmed by amplifying the two regions spanning the upstream and downstream insertion borders using PCR followed by Sanger sequencing (primer binding locations depicted in **Figure 7A**). The genotyping strategy is essentially as described for deletion allele generation via DMS cassette insertion in Norris et al. (2015).^54^

The forward and reverse primers used to amplify the upstream insertion region were GAAGTTTCCAAATGGTCGTAGAAC (located within the *nlg-1* genomic promoter region) and CGAGAAGCATTGAACACCATAAC (located within GFP in the selection cassette) respectively.

The forward and reverse primers used to amplify the downstream insertion region were TTCCTCGTGCTTTACGGTATCG (located within the Neomycin resistance gene) and GGTAGCTTGATTCGCCTTCTAT (located downstream of the *nlg-1* genomic coding region) respectively.

Following cassette excision via injection of cre-recombinase the *nlg-1* promoter (GAAGTTTCCAAATGGTCGTAGAAC) and *nlg-1* downstream (GGTAGCTTGATTCGCCTTCTAT) primers were used to amplify and confirm error free insertion of the *AID::GFP sequence* at the *nlg-1* locus via Sanger sequencing (**Figure 7**).

#### Auxin administration

Auxin administration was performed as previously described.^134^ Auxin treatment was performed by transferring animals to bacteria-seeded plates containing auxin. Auxin indole-3-acetic acid (IAA) (Thermo Fisher, Alfa Aesar™ #A1055614) was dissolved in ethanol to create a 400 mM stock solution. The stock solution was stored at 4°C for up to one month. Auxin was diluted into the molten NGM agar (cooled to ∼50°C before Auxin addition) before pouring plates. Auxin plates were seeded with 50 µl of OP50 liquid culture 12-24 hours before use. For continuous exposure groups, animals were age synchronized (as described above) for behavioral testing on auxin plates and tested at 96 hours old. For developmental exposure animals were age synchronized and reared on auxin plates for 48 hours before being transferred to standard OP50 seeded NGM plates, and then tested 48 hours later (96 hours old). For adult auxin treatment groups animals were age synchronized on standard OP50 seeded NGM plates and reared for 48 hours before being transferred to auxin plates and then tested 48 hours later (96 hours old).

#### Confocal imaging

Adult animals were anesthetised on glass microscope slides in 5 mM Levamisole and 150mM BDM (2,3-butanedione monoxime) dissolved in M9 buffer and covered with a 1.5 coverslip. A Leica SP8 white light laser confocal microscope and 60× oil immersion lens was used for imaging. Step size was 0.3 µm. GFP was excited using a 488 nm wavelength laser with emitted light collected through a 493–582 nm bandpass filter. YFP was excited using a 506 nm wavelength laser with emitted light collected through a 511-600 nm bandpass filter. mRuby was excited using a 587 nm wavelength laser with light collected through a 592-779 nm bandpass filter. Final figures were generated using ImageJ (National Institutes of Health, Bethesda, MD).^176^

## Acknowledgements

We would like to thank Dr. Evan L. Ardiel for useful comments and discussion regarding the manuscript. We would like to thank Dr. John Calarco, Dr. Erik Jorgensen, and Dr. Don Moerman and their labs for sharing their constructs and protocols or making them publicly available. We thank the *C. elegans* knockout consortium for several alleles, as well as the National Bioresource Project and the *C. elegans* Genetics Center (NIH Office of Research Infrastructure Programs, P40623 OD010440) for providing strains. We would like to thank Lexis Kepler and Anna Willms for assistance with behavioral and cloning experiments. We would like to thank Christine R. Ackerley for useful advice and discussions regarding figure design. We would also like to thank the *Caenorhabditis* Genetic Center and the National BioResource Project of Japan for strains.

## Competing interests

The authors declare no competing interests.

## Funding

This work was supported by a Canadian Institutes of Health Research Doctoral Research Award to TAM, Simons Foundation Autism Research Initiative (#205081) and Autism Speaks grants (#1975) to JBR, a SFARI award (#573845) to KH (PI), PP and CHR (co-PIs), and a Canadian Institutes of Health Research project grant (grant #CIHR MOP 130287) to CHR.

## Data availability

All data generated in this work is available free online in its raw and processed forms at (https://doi.org/10.5683/SP2/FJWIL8). All analysis code is freely available at (https://github.com/PavlidisLab/McDiarmid-etal-2019_Multi-Worm-Tracker-analysis). All strains and reagents are available from the corresponding author upon request.

## Author Contributions

TAM and CHR devised the concept of the study. TAM, MB, KH, PP, and CHR conceived the ASD gene selection, ortholog identification, and phenotypic characterization pipeline. MB, TAM, and PP wrote the analysis code. TAM and KM devised the AID modification to the Calarco CRISPR-Cas9 genome editing strategy, designed and built constructs. TAM, EAM, and FM designed reagents, performed cloning into expression vectors, prepared constructs for injection, and carried out genotype confirmation of transgenic animals. TAM, GPM, JBR, and JL generated the transgenic lines. TAM, GPM, JBR, and JL performed genotyping, genetic crosses, strain maintenance, and confocal imaging for the neuroligin strains, and designed and ran the relevant behavioral experiments. TAM and CHR designed the short-term habituation behavioral paradigm. TAM and MB made the figures. TAM drafted the manuscript, TAM, MB, JBR, KM, KH, PP, and CHR edited and co-wrote the final manuscript.

## Supplementary Figures

**Figure collection S1, Related to Figure 2. Reverse genetic screens.** All plots illustrate the sample mean distance of each genotype group from wild-type. Strains outside the 95% confidence interval of the wild-type distribution are labeled and colored blue. Only a maximum of ten strains are labeled in each direction per feature to prevent over plotting.

**Figure collection S2, Related to Figure 2. Phenotypic profiles.** For all plots bars represent directional t-statistics from an unpaired t-test comparing the indicated mutant to wild-type for each phenotypic feature listed across the x-axis. Color coding reflects feature classification.

**Figure collection S3, Related to Figure 4. Phenomic Heatmaps.** Phenomic heat maps summarizing the phenotypic profiles of 87 strains harbouring a mutation in an ortholog of an ASD-associated gene. Cells represent directional t-statistics from comparisons to wild-type controls. T-statistics are shown unclipped and at various clippings (t clipped at ±10, ±20, etc.). On select indicated heat maps, only cells significant at FDR < 0.1 are colored for ease of interpretation. The heat maps are interactive allowing for more detailed inspection of selected observations. Absolute t-statics values are clipped at 3.0, 10.0 and 20.0 in the last three figures.

**Figure collection S4, Related to Figure 4. Pvclust dendrograms.** Dendrograms depict hierarchical clustering of strains based on similarity in their phenotypic profiles. The Student’s T-statistic was used as a numerical score to represent the difference between wild-type and mutant animals for each phenotypic feature; this created a numerical profile of phenotypic features for further analysis. Average-linkage hierarchical clustering was performed with pvclust using correlation as the distance measure, and 50,000 rounds of bootstrapping. Clustering was performed on all features as well as the morphology features only and sensory and learning features only.

**Figure S5, Related to Figure 5.**
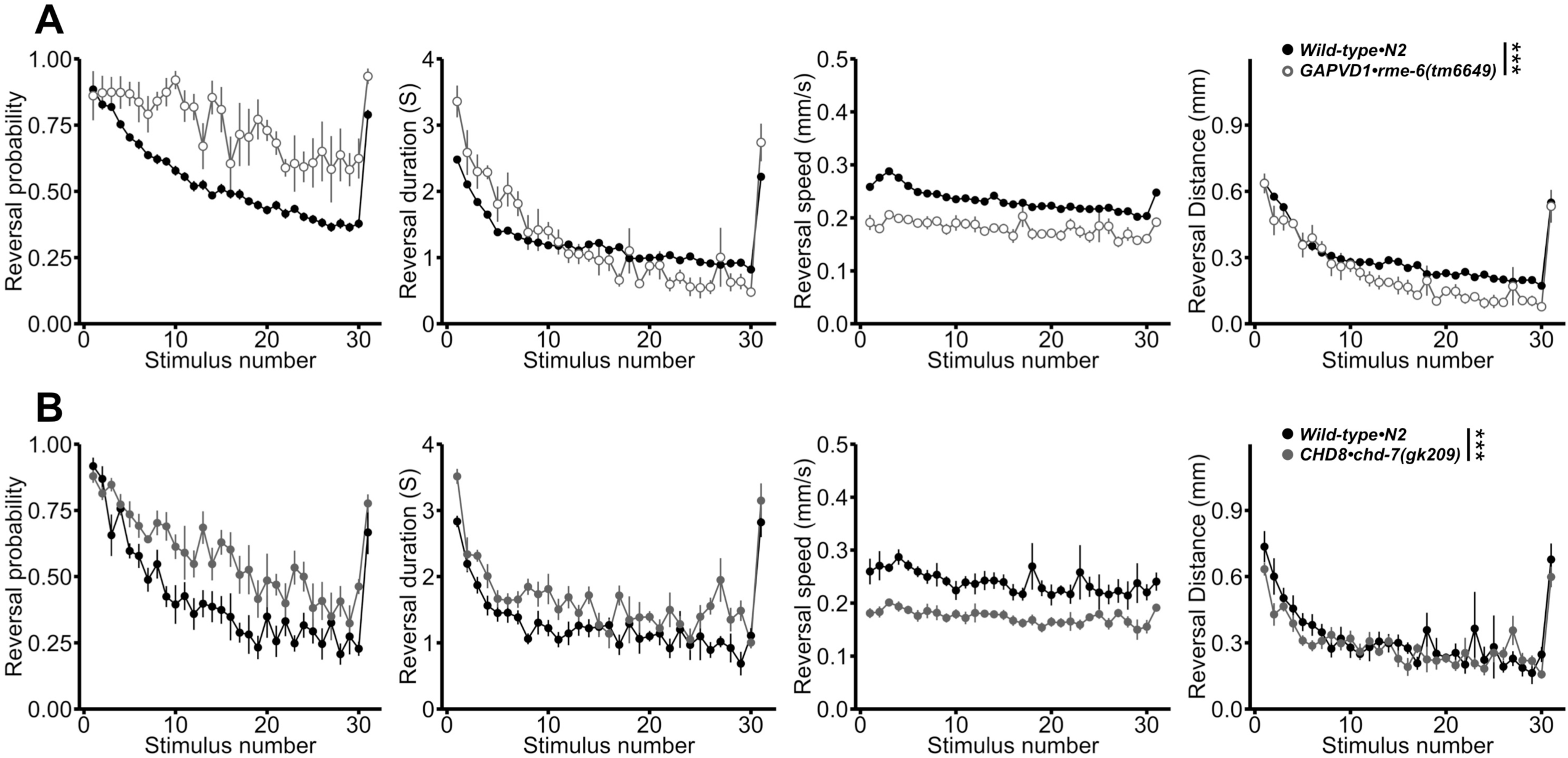
Second alleles of *GAPVD1•rme-6(tm6649)* and *CHD8•chd-7(gk209)* also display increased initial reversal response duration and impaired habituation of response probability. **A)** Sensory and learning phenotypic profile for *GAPVD1•rme-6(tm6659)* and **B)** *CHD8•chd-7(gk209)* mutants. Data are shown as mean±s.e.m using the number of plates as n. ***p<0.001, binomial logistic regression followed by Tukey’s HSD criterion was used to determine significance of the habituated level (proportion reversing at tap 30) for each pair of strains, n.s., not significant.

